# Structural basis for human TRPC5 channel inhibition by two distinct inhibitors

**DOI:** 10.1101/2020.04.21.052910

**Authors:** Kangcheng Song, Miao Wei, Wenjun Guo, Yunlu Kang, Jing-Xiang Wu, Lei Chen

## Abstract

TRPC5 channel is a non-selective cation channel that participates diverse physiological processes. Human TRPC5 inhibitors show promise in the treatment of anxiety disorder, depression and kidney disease. Despite the high relevance of TRPC5 to human health, its inhibitor binding pockets have not been fully characterized due to the lack of structural information, which greatly hinders structure-based drug discovery. Here we show cryo-EM structures of human TRPC5 in complex with two distinct inhibitors, namely clemizole and HC-070, to the resolution of 2.7 Å. Based on the high-quality cryo-EM maps, we uncover the different binding pockets and detailed binding modes for these two inhibitors. Clemizole binds inside the voltage sensor-like domain of each subunit, while HC-070 binds close to the ion channel pore and is wedged between adjacent subunits. Both of them exert the inhibitory function by stabilizing the ion channel in a closed state. These structures provide templates for further design and optimization of inhibitors targeting human TRPC5.

## Introduction

The mammalian transient receptor potential canonical (TRPC) channels are Ca^2+^-permeable nonselective cation channels that belong to the transient receptor potential (TRP) channel superfamily (*1*). Among all the subfamilies of TRP channels, TRPC share the closest homology to the Drosophila TRP channel, the first TRP channel cloned (*2*). The TRPC subfamily has seven members in mammals, TRPC1 to TRPC7, while TRPC2 is a pseudogene in human. According to the primary amino acid sequences, TRPC channels can be divided into TRPC1, TRPC2, TRPC3/6/7 and TRPC4/5 subgroups (*3*). Among them, TRPC5 is mainly expressed in the brain, and in liver, kidney, testis and pancreas to a lesser extent (*4-6*). It can form homotetrameric channels or heterotetrameric channels with TRPC4 and/or TRPC1 (*7, 8*). Upon activation, TRPC5 leads to depolarization of cell membrane and increase of intracellular Ca^2+^ level ([Ca^2+^]_i_). TRPC5 mediates diverse physiological processes and is implicated in many disease conditions such as fear, anxiety, depression and progressive kidney disease (*9, 10*).

Physiologically, TRPC5 channel is regulated by several mechanisms, including cell surface receptors, calcium stores, redox status and several cations. TRPC5 can be activated by receptor activated G_q/11_-PLC pathway upon the dissociation of Na^+^/H^+^ exchanger regulatory factor (NHERF) and activation of IP_3_ receptors (*11-13*), or through the G_i/o_ pathway (*14*). By interacting with STIM1, TRPC5 functions as a store-operated channel (*15, 16*). Extracellular application of reduced thioredoxin or reducing agents enhances TRPC5 activity (*17*). TRPC5 is also regulated by [Ca^2+^]_i_ in a concentration-dependent bell-shaped manner (*15, 18-20*) and extracellular Ca^2+^ ([Ca^2+^]_o_) at concentration higher than 5 mM robustly activates TRPC5 (*11, 15*).

Several available pharmacological tool compounds permit the identification of TRPC5 as putative drug targets for certain diseases (*21-23*). (-)-englerin A (EA), a natural product obtained from *Phyllanthus engleri*, can selectively activate TRPC4/5 and therefore inhibit tumor growth (*24, 25*), suggesting activation of TRPC4/5 might be a strategy to cure certain types of cancer. In contrast, TRPC5 inhibitors show promise for the treatment of central nervous system (CNS) diseases and focal segmental glomerulosclerosis (FSGS) in animal models (*26-28*). A TRPC4/5 inhibitor developed by Hydra and Boehringer Ingelheim is in clinical trial for the treatment of anxiety disorder and depression (*29*). Another TRPC5 inhibitor GFB-887 developed by Goldfinch Bio is currently in phase 1 clinical trial for the treatment of kidney disease (NCT number: NCT03970122).

Among TRPC5 inhibitors, clemizole (CMZ) and HC-070 represent two classes with distinct chemical structures. CMZ is a benzimidazole-derived H1 antagonist that can inhibit TRPC5. The same class of compounds includes M084 and AC1903 (*28, 30, 31*). In contrast, HC-070 is a methylxanthine derivative which inhibits TRPC4/5 with high potency (*27*). Another hTRPC5 inhibitor HC-608 and activator AM237 belong to this class (*27, 32*). The distinct sizes, shapes and polarities between CMZ and HC-070 type inhibitors suggest they bind hTRPC5 at different pockets which are still elusive due to the lack of structural information. This greatly impede further structural-based compound optimizations. To understand the inhibitory mechanism of these two distinct inhibitors against human TRPC5 (hTRPC5), we embarked structural studies of hTRPC5 in complex with CMZ or HC-070. Our high resolution cryo-EM maps are sufficient to identify their binding pockets and will aid further drug development.

## Results

### Structure of human TRPC5 in complex with inhibitors

To reveal the binding sites of CMZ and HC-070 on hTRPC5, we expressed hTRPC5 in HEK293F cells for structural studies. Similar to the mouse TRPC5 counterpart (mTRPC5) (*33*), C-terminal truncation of hTRPC5 generated a construct (hTRPC5_1-764_) that showed higher expression level of the tetramer in comparison with the full length wild type hTRPC5 (Fig. S1A). Moreover, we found hTRPC5_1-764_ retained the pharmacological properties of full length hTRPC5 such as activation by EA (Fig. S1, B to E). Therefore, we used hTRPC5_1-764_ protein for structural studies. Tetrameric hTRPC5_1-764_ channel was purified in detergent micelles (Fig. S1F and G). To obtain the CMZ-bound and the HC-070-bound TRPC5 structures, CMZ or HC-070 was added to the concentrated hTRPC5_1-764_ protein for cryo-EM sample preparation. We obtained maps of hTRPC5 in CMZ-bound state and HC-070-bound state to the resolution of 2.7 Å (Fig. 1, Figs. S2 to S5; table S1). The map qualities were sufficient for model building and ligand assignment (Fig. S3, D and E, and Fig. S5, D and E).

**Fig. 1.**
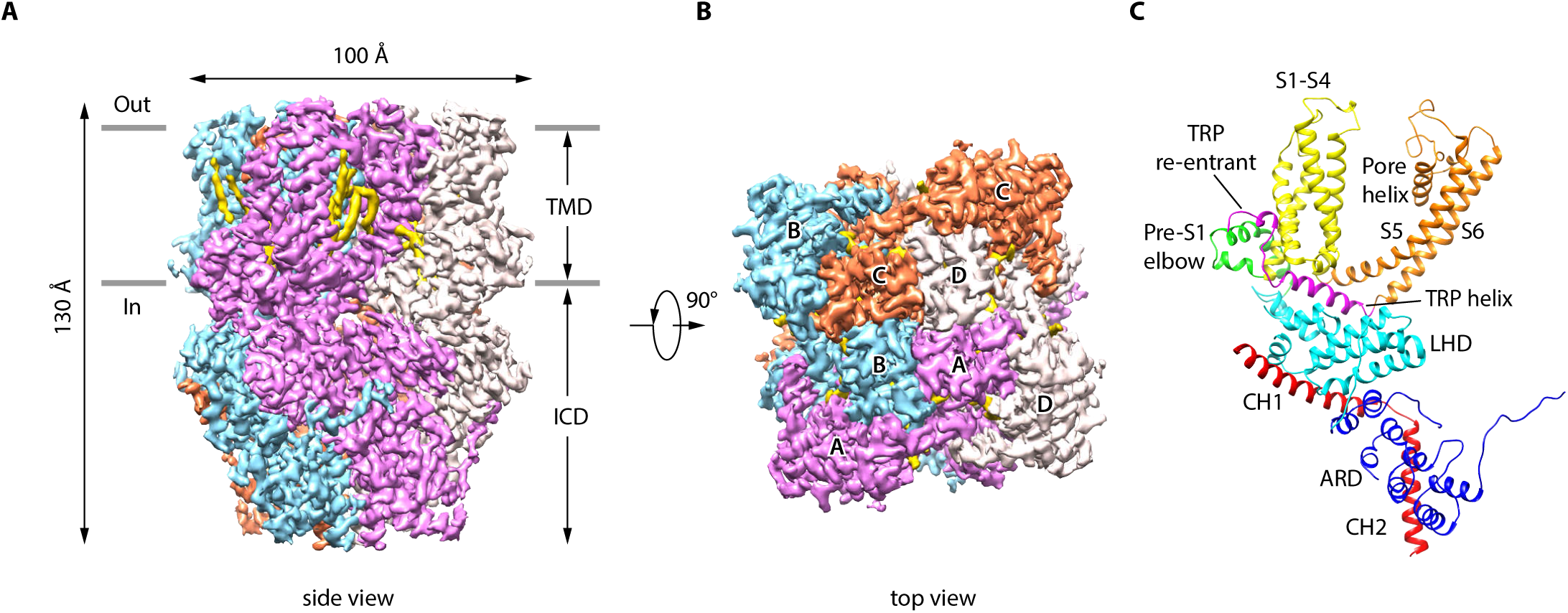
Overall structure of human TRPC5. (A, B) Cryo-EM density maps of CMZ-bound hTRPC5 shown in side view (A) and top view (B). Subunit A, B, C, and D are colored in purple, light blue, orange and gray, respectively. Lipids are colored in gold. The approximate boundary of the cell membrane is indicated by gray lines. TMD, transmembrane domain; ICD, intracellular cytosolic domain. (C) Ribbon diagram of a single subunit with secondary structure elements represented in different colors. ARD, ankyrin repeats domain; LHD, linker-helix domain; CH1, C-terminal helix 1; CH2, C-terminal helix 2.

The overall architecture of inhibitor-bound hTRPC5 is similar to the structure of mTRPC5 in apo state (*33*), occupying 100 Å × 100 Å × 130 Å in three-dimensional space (Fig. 1). The four-fold symmetric channel has two layer architecture, composed of the intracellular cytosolic domain (ICD) layer and the transmembrane domain (TMD) layer (Fig. 1). In ICD layer, the N-terminal ankyrin repeats domain (ARD) is below the linker-helix domain (LHD) region (Fig .1F). The TRP helix mediates interactions with TMD. The C-terminal helix 1 (CH1) and the C-terminal helix 2 (CH2) fold back to interact with LHD and ARD (Fig. 1F). In TMD layer, the pre-S1 elbow, the voltage sensor-like domain (VSLD, S1-S4), and the pore domain (S5-S6) assemble in a domain-swapped fashion. Densities of several putative lipid molecules were readily observed in the TMD layer, most of which are in the grooves between VSLD and the pore domains (Fig. 1, A and B).

### The CMZ binding site

By comparing the cryo-EM map of hTRPC5 in complex with CMZ, HC-070 and apo mTRPC5, we unambiguously located the binding site of CMZ and HC-070 (Fig. S3 and S5). A non-protein density located within the pocket surrounded by S1-S4 and TRP re-entrant loop of VSLD was observed only in the presence of CMZ and its size and shape match those of CMZ compound (Fig. 2, A and B, and Fig. S3E). Therefore, we assign this density as CMZ. Several residues in hTRPC5 are in close proximity with CMZ (Fig. 2C), including Y374 on S1, F414, G417 and E418 on S2, D439, M442, N443 and Y446 on S3, R492 and S495 on S4 and P659 on TRP re-entrant loop (Fig. 2, C and D). Among them, F414 forms a π-π stacking interaction with the chlorophenyl ring of CMZ. Side chain of Y374, M442, Y446 and P659 show hydrophobic interactions with CMZ. Residues related to CMZ binding are highly conserved between TRPC4 and TRPC5 which suggest the same binding pocket both in TRPC5 and TRPC4 (Fig. S6).

**Fig. 2.**
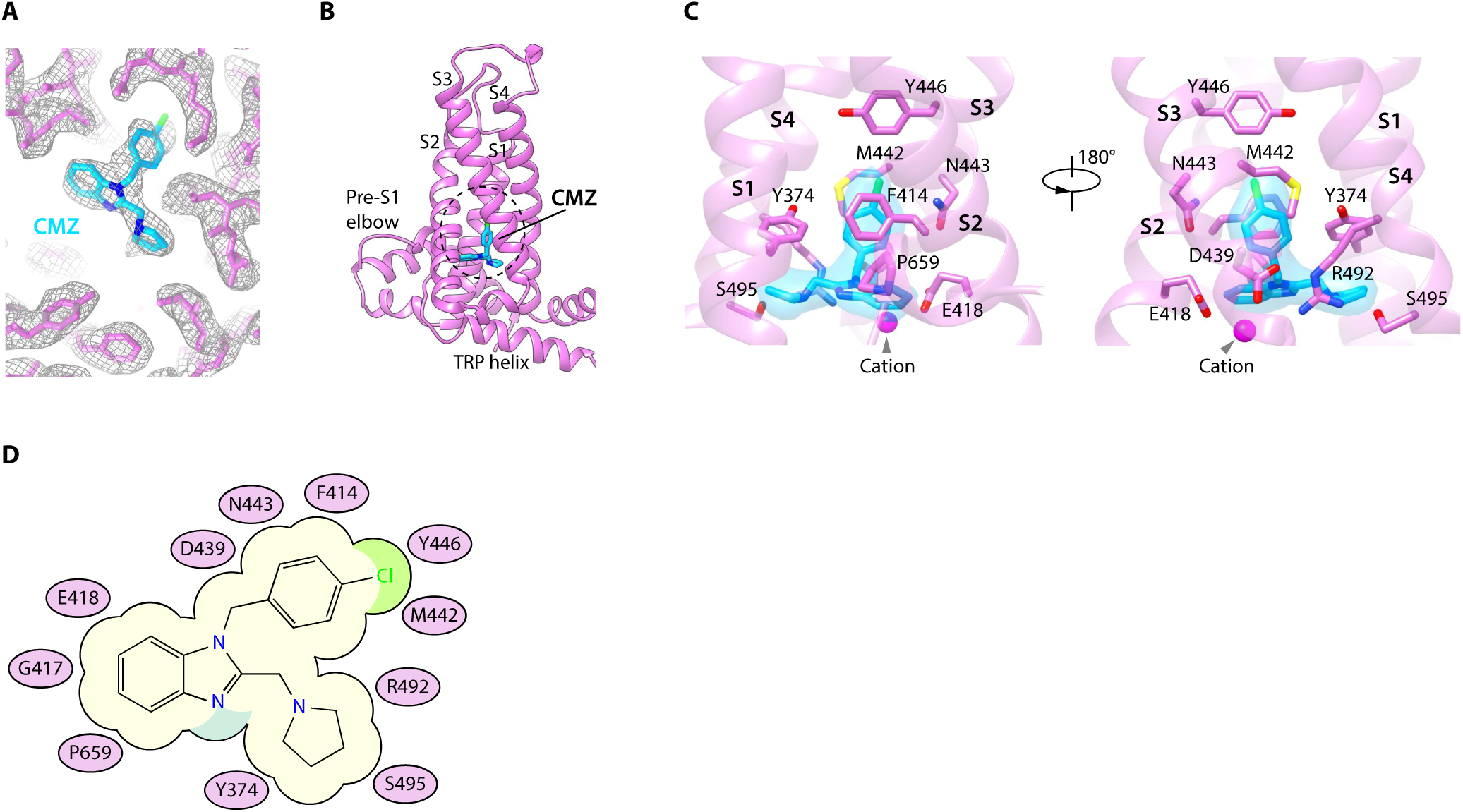
CMZ binding site in hTRPC5. (A) Densities of CMZ and nearby residues. The map is contoured at 4.6 σ (gray mesh). CMZ and hTRPC5 are shown in sticks, colored in sky blue and purple, respectively. (B) Overview of the CMZ binding site in hTRPC5. The dashed region demarcates the binding pocket of CMZ. (C) Close-up view of the CMZ binding site. CMZ is shown as transparent surface superimposed with sticks. Side chains of residues that interact with CMZ are shown as sticks. A cation near CMZ is shown as a magenta sphere. (D) Cartoon representation of the interactions between CMZ and hTRPC5. S1-S4 of subunit A are represented as purple ovals. Residues that interact with CMZ are labeled inside the ovals.

### The HC-070 binding site

A non-protein density surrounded by the S5, pore helix from one subunit and S6 of the adjacent subunit was observed in the presence of HC-070 (Fig. 3, A and B, and Fig. S5E). Moreover, the size and shape of this density perfectly match those of HC-070, suggesting this density might represent HC-070 compound. HC-070 is a clover-shaped molecule with a hydroxypropyl tail, a chlorophenyl ring and a chlorophenoxy ring which extend out from the methylxanthine core. hTRPC5 makes extensive interactions with HC-070 (Fig. 3, C and D). The side chain of F576 on pore helix forms π-stacking interaction with the methylxanthine core of HC-070. R557 on the loop linking S5 and pore helix (Linker), Q573 and W577 on pore helix form polar contacts with the hydroxypropyl tail of HC-070. F599, A602 and T603 on S6 from the adjacent subunit also stabilize the hydroxypropyl tail. F569 and L572 interact hydrophobically with the chlorophenyl ring of HC-070. The chlorophenoxy branch only weakly interacts with C525 and this might account for its weaker density and higher flexibility (Fig. S5E). Residues involved in HC-070 binding are absolutely conserved between hTRPC4 and hTRPC5 but not in TRPC3/6/7 (Fig. S6), in agreement with the fact that HC-070 is much more potent toward TRPC4/5 than toward TRPC3/6/7 (*27*).

**Fig. 3.**
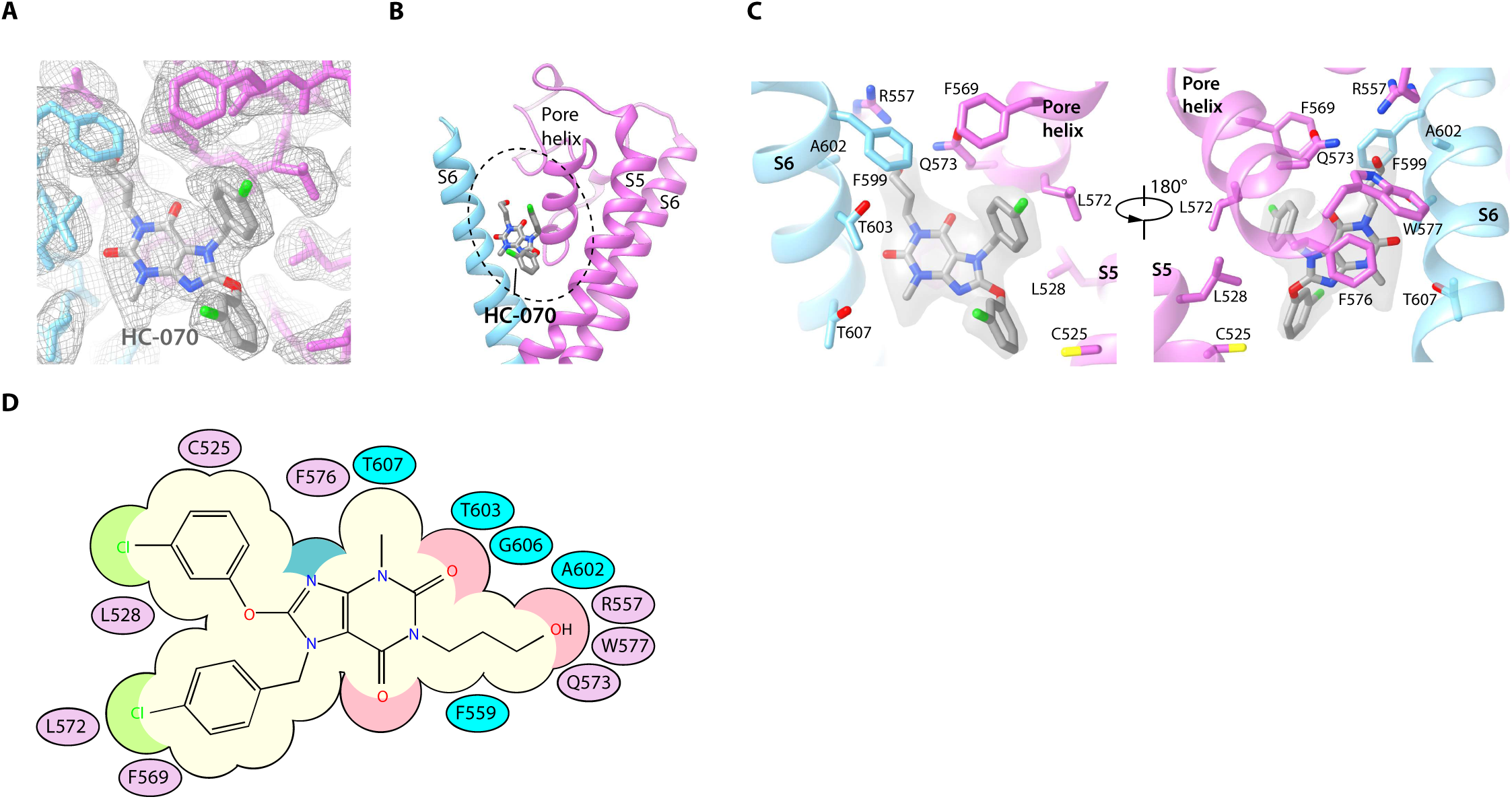
HC-070 binding site in hTRPC5. (A) Densities within the HC-070 binding site. The map is contoured at 3 σ (gray mesh). HC-070 and hTRPC5 are shown in sticks, with HC-070 colored in gray. Subunits A and B are colored in purple and light blue, respectively. (B) Overview of the HC-070 binding site in hTRPC5. The dashed region marks the binding pocket of HC-070. (C) Close-up view of the HC-070 binding site. HC-070 is shown as transparent surface superimposed with stick-model. Side chains of residues that interact with HC-070 are shown as sticks. (D) Cartoon representation of the interactions between HC-070 and hTRPC5. S5 of Subunit A and S6 of subunit B are represented as purple and light blue ovals, respectively. Residues that interact with HC-070 are labeled inside the ovals.

### Ion conduction pore and non-protein densities in cryo-EM maps

In TMD, pore helixes, pore loops and S6 of four protomers form the transmembrane pathway for ion permeation. Calculated pore profiles showed that the constriction is formed by the side chains of I621, N625 and Q629, which is the lower gate of hTRPC5 (Fig. 4, A and B). The smallest radius at the gate are below 1 Å, indicating the channel is in a non-conductive closed state. This is in agreement with the inhibitory function of CMZ and HC-070. A putative cation density inside VSLD, surrounded by E418, E421 on S2 and N436, D439 on S3, was observed in both CMZ and HC-070 maps (Fig. 4C). Strikingly, this putative cation is in close proximity to CMZ in CMZ-bound hTRPC5 (Fig. 2C). Similar density was previously observed in mTRPC4, mTRPC5, TRPM2, TRPM4 and TRPM8 (*33-37*). The identity and function of this putative ion still await further investigation. In ICD, another extra density is found in the ankyrin repeat 4 (AR4)-linker helix (LH1) region (Fig.4D). The ion is coordinated by the side chains of H178 and C182 on AR4-LH1 loop and C184 and C187 on LH1 helix (Fig. 4D). Based on the chemical environments for ion coordination, this ion binding site is likely a HC3 type zinc binding site, as observed previously in a bacteria glucokinase (*38*). Because the zinc ion is from endogenous source and co-purified with hTRPC5, we speculate this is a high affinity zinc binding site that constitutively chelates zinc. The residues coordinating putative zinc ions are highly conserved in TRPC ion channels (Fig. S6), but the physiological function of this site still awaits further investigation.

**Fig. 4.**
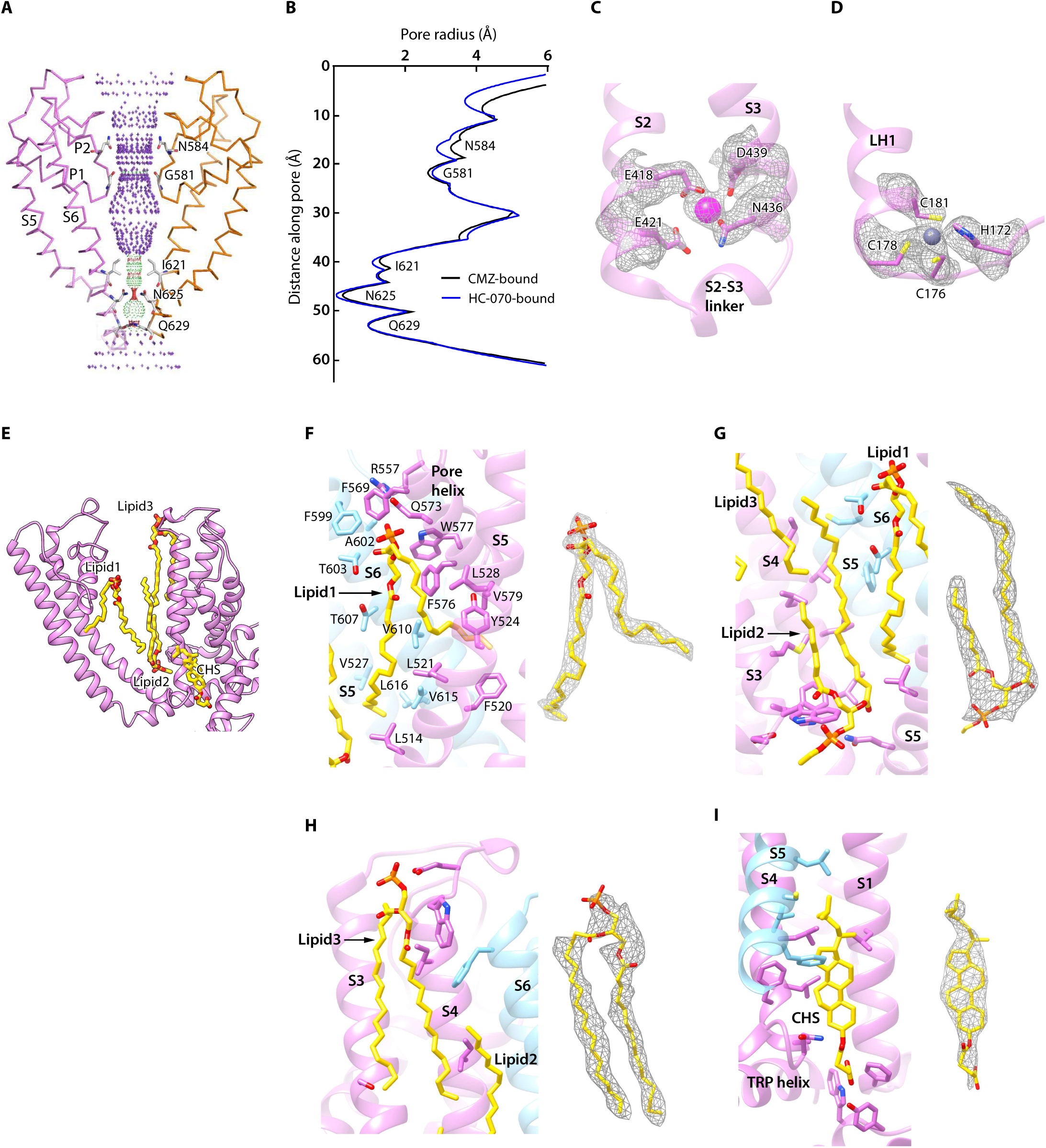
Ion conduction pore and other non-protein densities in hTRPC5 maps. (A) Side view of the pore region of non-conductive hTRPC5 (CMZ-bound). Subunit A and subunit C are shown as ribbon and colored the same as in Fig.1. Subunits B and D are omitted for clarity. Ion conduction pathway along the pore is shown as dots with key residues labeled, calculated by HOLE. Purple, green and red dots define pore radii of >2.8 Å, 1.4–2.8 Å and <1.4 Å, respectively. (B) Calculated pore radius of CMZ-bound hTRPC5 and HC070-bound hTRPC5 are shown vertically. (C) Close up view of the putative cation binding site in TMD. Densities of the cation and the related residues are contoured at 3.2σ and shown as grey mesh. The side chain of the residues that interact with the cation are shown as sticks. (D) Close up view of the putative zinc binding site in hTRPC5. Densities of the zinc ion and interacting residues are contoured at 3.1σ and shown as grey mesh. (E) Other modeled non-protein densities in TMD in CMZ-bound hTRPC5 are shown as sticks and colored the same as in Fig. 1. (F-I) Close-up views of lipid1 (F), lipid2 (G), lipid3 (H) and CHS (I) binding sites. Subunit A and B are colored in purple and light blue, respectively. Residues that interact with lipids are shown as sticks. Insets show cryo-EM densities of these lipids, contoured at 5σ, 4.7σ, 3σ, 5.1σ, respectively.

In the 2.7 Å cryo-EM maps of hTRPC5, we observed several putative lipid molecules in the transmembrane domain (Fig. 1, A and B and Fig. 4E). Lipid 1 locates near the S5 and pore helix from one subunit and S6 from the adjacent subunit in the CMZ-bound hTRPC5. Its polar head group faces the extracellular side and forms polar contacts with R557 on Linker, Q573 and W577 on pore helix, suggesting its outer leaflet localization (Fig. 4F). Its two hydrophobic tails interact with several hydrophobic residues F576, V579 on pore helix, L514, F520, L521, Y524 on S5 and V610, V614, L616 on S6. Similar densities were observed in mTRPC4 and mTRPC5 (*33, 34*), suggesting the important function of lipid 1. Interestingly, in the map of HC-070-bound hTRPC5, the head group of lipid 1 is replaced by HC-070, while the tails of lipid 1 are still in similar position (Fig. S5F). Lipid 2 and lipid 3 are observed in both CMZ-bound hTRPC5 and HC-070-bound hTRPC5 maps. Lipid 2 localizes in the inner leaflet and is bound by S3, S4 and S5 from one subunit and S5, S6 from adjacent subunit (Fig. 4G). Lipid 3 is in the outer leaflet of membrane and bound by S3 and S4 (Fig. 4H). There is one putative cholesteryl hemisuccinate (CHS) molecule which is sandwiched by S1, S4 from one subunit and S5 from adjacent subunit (Fig. 4I). Densities similar to the putative CHS were observed previously in corresponding positions in TRPM4 and TRPCs (*33, 34, 36, 39*). Moreover, this putative CHS binding site was reported to be important for the activation of mTRPC5 (*33*), suggesting its potential role in channel modulation.

### Structure comparisons between available TRPC5 structures

The structure of truncated mTRPC5 in the apo state was solved to the resolution of 2.8 Å previously (*33*). The constructs of truncated hTRPC5 and mTRPC5 used for cryo-EM studies share over 99% sequence identity (Fig. S6). In agreement with the high sequence conservation, the overall structure of hTRPC5 and mTRPC5 (PDB ID: 6AEI) are highly similar, with root-mean-square deviation (RMSD) of 0.6 Å (0.635 Å between CMZ-bound hTRPC5 and 6AEI; 0.617 Å between HC070-bound hTRPC5 and 6AEI), although there are several structural variations in details (Fig. 5).

**Fig. 5.**
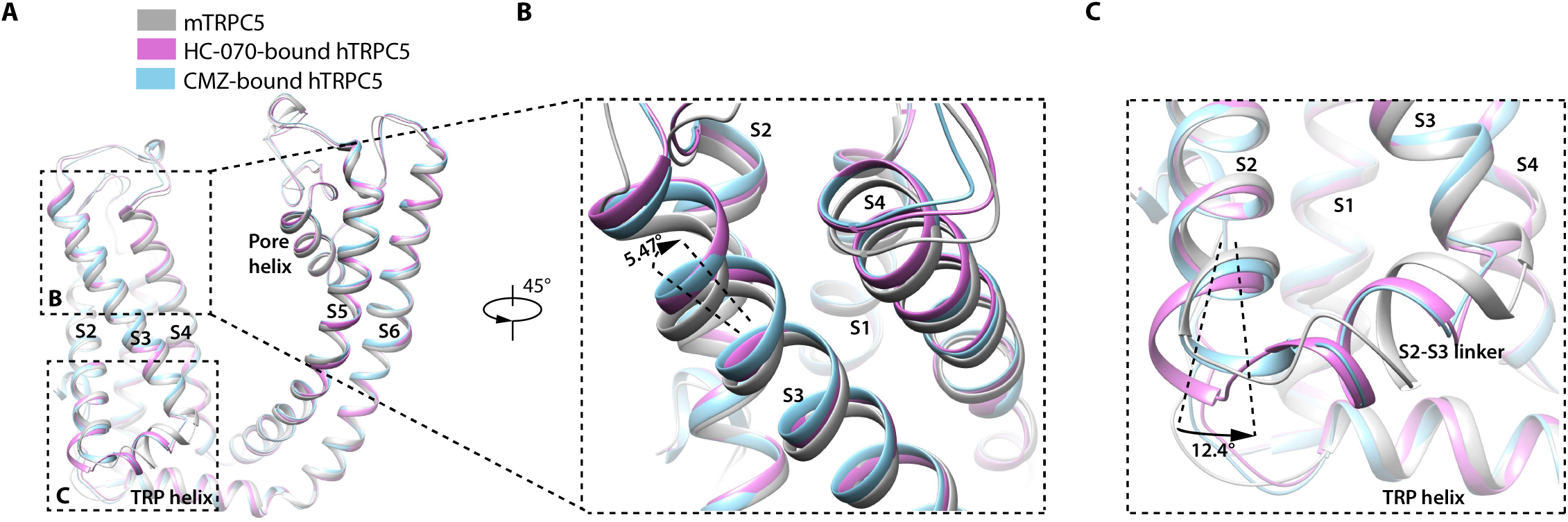
Structure comparison between CMZ-bound TRPC5, HC-070-bound TRPC5 and apo mTRPC5 in TMD. (A) The overall conformational variation in TMD between these three structures. All the structures are shown as cartoon, with apo mTRPC5 colored in grey, CMZ-bound TRPC5 in light blue and HC-070-bound TRPC5 in purple. (B) Close up view of the conformational change of S3 in both CMZ-bound TRPC5 and HC-070-bound TRPC5 structures compared to the apo mTRPC5 structure. The tilt angle of S3 (444-457 aa) was measured when these TRPC5 structures are aligned. (C) Close up view of the structure comparison of S2 between these three structures. The tilt angle of helix (419-425 aa) was measured when the structure of CMZ-bound TRPC5 and HC-070-bound TRPC5 are aligned.

The CMZ bound and the HC-070 bound structures have some subtle conformational change compared with ligand-free mTRPC5 in TMD (Fig. 5A). S3 segments on the extracellular side in CMZ and HC-070 bound structures tilt for 5.47 degree compared to the ligand-free mTRPC5 (Fig. 5B). Moreover, the S2-S3 linker helixes move half a helix away from the central axis of the channel in inhibitor-bound structures (Fig. 5C). These subtle variations may be due to different sample preparation procedures between hTRPC5 (in GDN detergent) and mTRPC5 (in PMAL-C8 amphipol). However, S2 of HC-070-bound hTRPC5 structure tilt 12.4 degree compared with CMZ-bound structure, probably due to inhibitor binding (Fig. 5C).

## Discussion

Thanks to the efforts from several groups (*40, 41*), three inhibitor binding pockets have been structurally identified in hTRPC5 and hTRPC6, namely inhibitor binding pocket (IBP) A-C (Fig. 6 and Fig. S7).

**Fig. 6.**
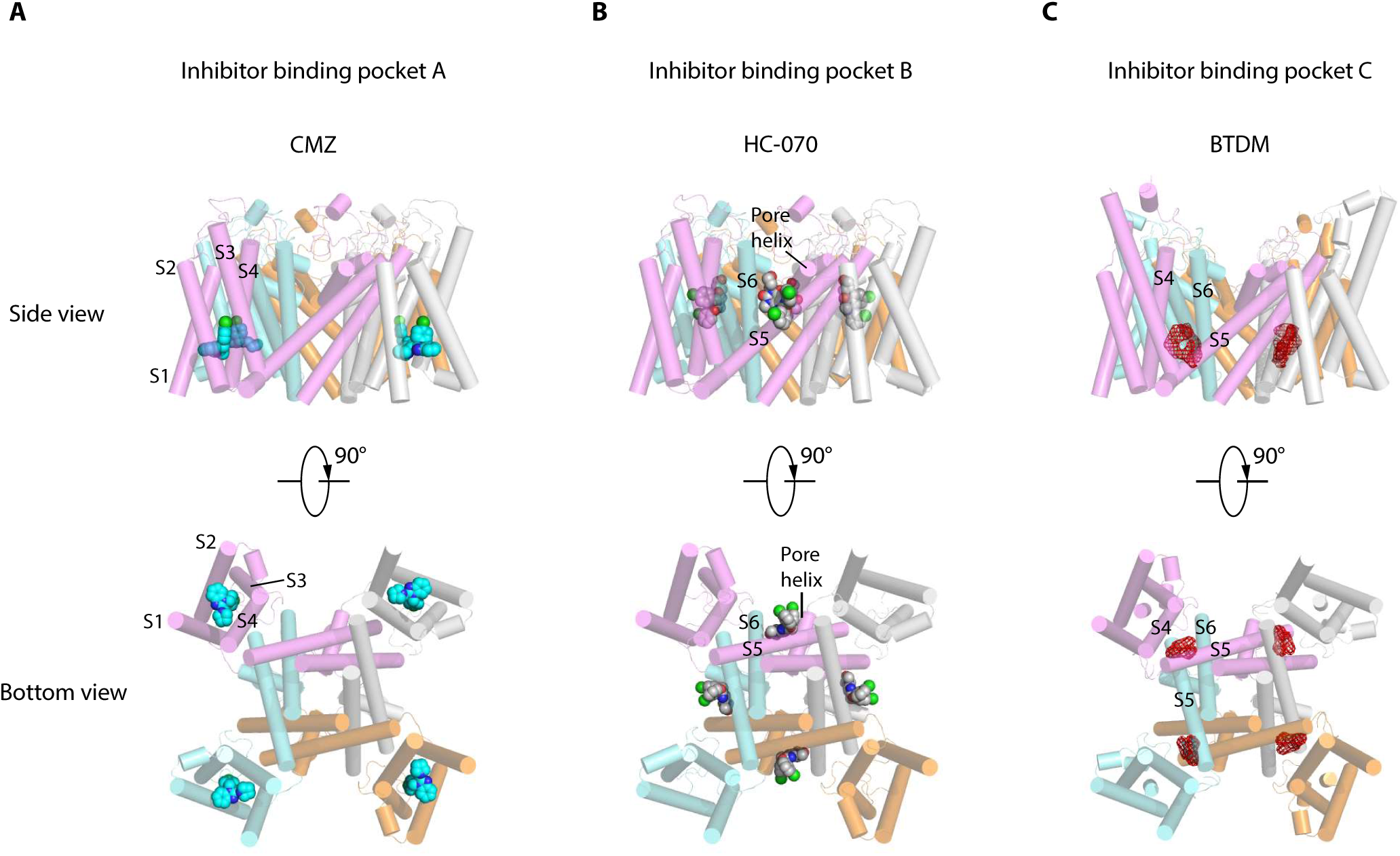
Inhibitor binding pockets of TRPCs in TMD. (A) CMZ binding pocket in hTRPC5 is shown in side view and bottom view. The TRPC5 structure is shown as cylinder with CMZ compound shown as spheres and colored in sky blue. Each subunit of TRPC5 is colored the same as Fig. 1. (B-C) HC-070 binding pocket in hTRPC5 structure and BTDM binding pocket in TRPC6 structure are shown in side view and bottom view. HC-070 is shown as spheres and colored in gray. The densities of BTDM are shown as red mesh.

IBP-A inside the VSLD is accessible from intracellular side (Fig. 6A). Inhibitors, such as CMZ, need to penetrate the cell membrane before binding at IBP-A. CMZ, M084 and AC1903 are benzimidazole derivatives which can inhibit TRPC5 with IC_50_ values at 1.0 - 1.3 μM, 8.2 μM, 14.7 μM, respectively (*28, 30, 31*). Their similar chemical structures suggest they probably share a common binding site at IBP-A (Fig. S8, A to C). CMZ are six-fold more potent against TRPC5 than TRPC4β (*31*). Residues in the IBP-A are highly conserved between TRPC5 and TRPC4, except P659 in TRPC5 is replaced by a threonine in TRPC4 (Fig. S6), which might be responsible for the observed six-fold selectivity of CMZ between TRPC4 and TRPC5. In addition, CMZ is ten-fold and 26-fold more potent against TRPC5 than TRPC3/6 and TRPC7, respectively (*31*). Residues E418, D439, M442, R492 are identical among TRPC4/5/3/6/7, whereas residues Y374 in TRPC5 are substituted by Phe, F414 by Met, N443 by Leu, Y446 by Phe and S495 by Tyr in TRPC3/6/7, which might be responsible for the lower potency of CMZ against TRPC3/6/7. Although VSLD has lost the voltage sensing function, it plays important structural and functional role in TRP channels. Based on the gating model of TRPV1, channel opening is associated with relative movements between VSLD and pore (*42, 43*) and thus, stabilization of VSLD in an inactive conformation might be a common scheme to inhibit channel opening. Indeed, AM-1473, a potent antagonist of hTRPC6 were found to bind at IBP-A in hTRPC6 as well (*44*). Moreover, IBP-A is the AMTB and TC-I 2014 binding sites in TRPM8 (*45*), 2-APB binding site in TRPV6 (*46*) (Fig. S7A).

IBP-B locates in the pore domain, sandwiched between subunit interface (Fig. 6B). HC-070 binds at IBP-B which is close to the extracellular side. Strikingly, IBP-B is partially occupied by the head group of a lipid molecule in the map of CMZ-bound hTRPC5 or apo mTRPC5 structures (Fig. S7B). Previous studies showed disruption of this lipid binding site in mTRPC5 by mutations, such as F576A and W577A, renders mTRPC5 inactive (*33*). In addition, mutation of W680 to alanine in hTRPC6, corresponding to W577A in hTRPC5, abolished OAG activation of hTRPC6 (*44*). Mutations of G709/G640 in hTRPC6/3, corresponding to G606 in hTRPC5 in IBP-B, blunted PLC-mediated activation of TRPC6/3, altered responses to DAGs of TRPC3, and also altered the pattern of photoactivation of TRPC6/3 by a photo-switchable DAG analogue, OptoDArG (*47*). These data collectively suggest this lipid binding pocket is important for the function of TRPC channels and ligands binding at IBP-B might affect TRPC channel gating. The binding of HC-070 at this site stabilizes hTRPC5 in a closed state and thus inhibits channel opening. The residues involved in HC-070 binding are absolutely identical between hTRPC5 and hTRPC4, but they share only 20% similarities between TRPC4/5 and TRPC3/6/7 (Fig. S6). F576 and W577 in the highly conserved LFW motif and G606 out of 13 residues in IBP-B are the identical residues between hTRPC5 and TRPC3/6/7, which explains the high selectivity of HC-070 for TRPC4/5 over TRPC3/6/7 (*27*). In addition, HC-070 shows inhibitory effect on heterotetrameric hTRPC1:C5 and hTRPC1:C4 with relatively lower potency compared to homotetrameric hTRPC4/5 (*27*), probably because of the lower sequence identity between TRPC4/5 and TRPC1 (46% according to the sequence alignment) (Fig. S6). HC-608, an HC-070 analogue, is more potent against hTRPC4/5 than HC-070 (*27*), probably due to the substitution of chloride with trifluoromethoxy group on the chlorophenoxy ring of HC-070 (Fig. S8, D and E). Unexpectedly, another HC-070 family compound, AM237, is a TRPC5 activator with EC_50_ around 15 - 20 nM (*32*). Compared with HC-608, AM237 has an additional chloride atom on the phenoxy ring (Fig. S8F). It is reported that AM237 activates homomeric hTRPC5 channel instead of the heterotetrameric TRPC1:C5 channel (*32*). However, AM237 inhibits EA-evoked activation of both homotetrameric and heterotetrameric TRPC5 channels (*32*), supporting that AM237 also binds at IBP-B which is on the interface between adjacent subunits. The structural mechanism of how AM237 activates hTRPC5 remains elusive. Recently, a hTRPC6 agonist AM-0883 was found to bind at IBP-B as well (Fig. S7B) (*44*), suggesting IBP-B is a common modulatory ligand binding site in TRPC channels.

IBP-C is between VSLD and pore domain, formed by S3, S4, S4-S5 linker from one subunit and S5-S6 from the adjacent subunit (Fig. 6C). BTDM, a high affinity inhibitor of hTRPC6 binds at IBP-C (Fig. 6C and Fig. S7C) (*39*). When BTDM is not supplied, IBP-C is occupied by a putative phospholipid (*44*). In TRPC4/5, mutations of G503/G504 to serine on S4-S5 linker lead to constitutive activation of the channels (*48*). In addition, vanilloid agonists resiniferatoxin (RTX) and capsaicin, vanilloid antagonist capsazepine, and an endogenous phosphatidylinositol bind at IBP-C on TRPV1 (Fig. S7C) (*42, 43*). These data together suggest IBP-C is important for the gating of both TRPC and TRPV channels.

In summary, our high resolution structures of hTRPC5 reported here have uncovered the binding pockets for two distinct classes of inhibitors together with their detailed binding mode. Further structural comparisons with previously reported TRP channel structures have revealed three common inhibitor binding pockets in TRPC channels. Our studies paved the way for further mechanistic studies and structure-based drug development.

## Materials and Methods

### FSEC

The cDNA of hTRPC5 and related mutants were all cloned into a home-made BacMam vector with N terminal GFP-MBP tag (*49*). The expression levels of various hTRPC5 constructs were screened by FSEC (*50*). Various constructs were transfected into HEK293F cells using PEI. After transfection for 40 h, cells were harvested by centrifuge at 5,000 rpm for 5 min. Then the cells were solubilized by 1% (w/v) LMNG, 0.1% (w/v) CHS in TBS buffer (20 mM Tris-HCl, pH 8.0 at 4 °C, and 150 mM NaCl) with 1 ug/ml aprotinin, 1 ug/ml leupeptin, 1 ug/ml pepstatin for 30 min at 4 °C. Cell supernatant were collected after centrifuge at 40,000 rpm for 30 min and loaded onto superpose 6 increase (GE healthcare) for FSEC analysis.

### FLIPR calcium assay

AD293 cells cultured on 10 cm dish (grown in Dulbecco’s Modified Eagle’s Medium + 10% FBS, 37 °C) at 100% confluence w ere digested with trypsin-EDTA (0.25%) (Gibco) and diluted for 3 fold with culture medium. Then the AD293 cells were seeded on black well, clear bottom, Poly-D-lysine coated 96-well plate (Costar) and cultured overnight. Then hTRPC5 constructs were transfected into AD293 cells (60%∼80% confluence) with PEI. About 24 hours after transfection, culture medium were discarded and cells were incubated with 50 μl per well of 4 μM Rhod-2/AM (AAT Bioquest) in TARODES’s buffer (10 mM HEPES, 150 mM NaCl, 4 mM KCl, 2 mM MgCl_2_, 2 mM CaCl_2_, 11 mM glucose, pH 7.4) supplemented with 0.02% Pluronic F-127 (Sigma) for 5 min in dark at room temperature. Then dye was removed and replaced with 50 μl of TARODE’s buffer per well. Calcium signals evoked by 256 nM EA (prepared in TARODES’s buffer) were detected using FLIPR-TETRA (Molecular Devices) with excitation/emission at 510–545/565–625 nm.

### Protein expression and purification

The BacMam expression system was established for large-scale expression of hTRPC5_1-764_ as previously reported (*40*). Briefly, hTRPC5_1-764_ was subcloned into a home-made BacMam vector with N terminal GFP-MBP tag. The Bacmid was then generated by transforming this construct into DH10Bac E. coli cells. Baculovirus was harvested about 7 days after transfecting bacmid into Sf9 cells cultured in Sf-900 III SFM medium (Gibco) at 27 °C. P2 baculovirus was produced using Bac-to-bac system. P2 virus was added to FreeStyle 293-F cells at a ratio of 1:12.5 (v/v) when the cells were grown to a density of 2.0×10 ^6^/ml in SMM 293T-I medium (Sino Biological Inc.) supplemented with 1% FBS under 5% CO2 in a shaker at 130 rpm at 37°C. After virus infection for 12 hours, 10 mM sodium butyrate was added and temperature was lowered to 30 °C. Cells were harvested 48 h post-infection and washed twice using TBS buffer. Cell pellets were collected and stored at −80 °C for further purification.

Cell pellets were resuspended in TBS buffer supplemented with 1% (w/v) LMNG, 0.1% (w/v) CHS, 1 mM dithiothreitol (DTT), 1 mM phenylmethanesulfonylfluoride (PMSF), and protease inhibitors, including 1 ug/ml aprotinin, 1 ug/ml leupeptin, 1 ug/ml pepstatin. The mixture was incubated at 4 °C for 1 h. After ultra-centrifugation at 40,000 rpm for 1 h in Ti45 rotor (Beckman), the supernatant was loaded onto 7 ml MBP resin. The resin was rinsed with wash buffer 1 (wash buffer 2 + 1mM ATP + 10 mM MgCl_2_) and wash buffer 2 (TBS + 40 μM glycol-diosgenin (GDN) + 0.005 mg/ml SPLE + 1 mM DTT) subsequently. Proteins were eluted with 80 mM maltose in wash buffer 2. The eluate was concentrated using a 100-kDa cut-off concentrator (Millipore) after digestion with H3C protease at 4 °C overnight. hTRPC5 protein was further purified by size exclusion chromatography (Superose-6, GE Healthcare) in buffer containing TBS, 40 μM GDN, 0.005 mg/ml SPLE, and 1 mM Tris (2-carboxyethyl) phosphine (TCEP). The peak fractions corresponding to tetrameric TRPC5 channel were collected and concentrated to A280 = 1.0 with estimated concentration of 0.7 mg/ml for sample preparation.

### Cryo-EM sample preparation and data collection

Purified protein was mixed with 50 μM HC-070, 100 μM Ca^2+^, or 500 μM Clemizole (CMZ) in TBS buffer containing 100 μM Ca^2+^, respectively. Aliquots of 2.5 μl mixture were placed on GO-grids as previously reported (*51*). Grids were blotted for 3 s (HC-070-bound) or 5 s (CMZ-bound) at 100% humidity and flash-frozen in liquid ethane cooled by liquid nitrogen using Vitrobot Mark I (FEI). Grids were then transferred to a Titan Krios (FEI) electron microscope that was equipped with a Gatan GIF Quantum energy filter and operated at 300 kV accelerating voltage. Image stacks were recorded on a Gatan K2 Summit direct detector in super-resolution counting mode using SerialEM at a nominal magnification of 165,000 × (HC-070-bound, calibrated pixel size of 0.821 Å/pixel) or 13,000 × (CMZ-bound, calibrated pixel size of 1.045 Å/pixel), with a defocus ranging from −1.5 to −2.0 μm. Each stack of 32 frames was exposed for 7.1s, with a total dose about 50 e^-^/ Å ^2^ and a dose rate of 8 e^-^/pixel/s on detector.

### Image processing

The CMZ-bound TRPC5 dataset and HC-070-bound TRPC5 dataset were processed with similar workflow. The collected image stacks were motion corrected by MotionCor2 (*52*). After motion correction, a total of 1,268/991 (CMZ/HC-070) good micrographs were manually selected, then GCTF (*53*) was used for CTF estimation. Following 354,312/274,189 particles were auto-picked using Gautomatch based on the projection of hTRPC6 map (EMDB: EMD-6856) for CMZ-bound or HC-070-bound hTRPC5, respectively. After two-rounds reference-free two-dimensional (2D) classification, a total of 109,369/114,211 (CMZ-bound/HC-070-bound) particles were selected for three-dimensional (3D) classification with previous hTRPC6 map (EMDB: EMD-6856) low pass filtered to 18 Å as the initial model using RELION-2.0 (*54*) with C1 symmetry. After 3D classification, classes containing 90,446/114,211 (CMZ-bound/HC-070-bound) particles with clearly visible α-helices in TMD were selected for further homogeneous refinements in cryoSPARC (*55*) using C4 symmetry. Reported resolutions are based on the gold-standard Fourier shell correlation (FSC) 0.143 criterion after correction of masking effect (*56*). The map was sharpened with a B factor determined by cryoSPARC automatically.

### Model building, refinement, and validation

The previously reported TRPC4 (PDB: 5Z96) was used as the starting model and docked into the HC-070-bound or CMZ-bound hTRPC5 maps in Chimera (*57*), respectively. The fitted model was manually adjusted in COOT (*58*), keeping the side chains of conserved residues and substituting non-conserved residues based on the sequence alignment between mTRPC5 and hTRPC5. The ligands models were generated using elbow (*59*) module in PHENIX (*60*). The ligands and phospholipids were manually docked into densities and refined using COOT. The models were further refined against the corresponding maps with PHENIX (*60*).

### Quantification and statistical analysis

The local resolution map was calculated using cryoSPARC. The pore radius was calculated using HOLE (*61*).

### Data availability

The density maps of hTRPC5 have been deposited to the Electron Microscopy Data Bank (EMDB) under the accession number: EMD-XXXX for HC-070-bound hTRPC5 and EMD-XXXX for CMZ-bound hTRPC5. Coordinates of atomic model have been deposited in the Protein Data Bank (PDB) under the accession number: XXXX for HC-070-bound hTRPC5 and XXXX for CMZ-bound hTRPC5. All other data are available from the corresponding authors upon reasonable request.

## Acknowledgement

The cDNAs of hTRPC5 were kindly provided by Xiaolin Zhang. Cryo-EM data collection was supported by Electron microscopy laboratory and Cryo-EM platform of Peking University with the assistance of Xuemei Li, Daqi Yu, Xia Pei, Bo Shao, Guopeng Wang, and Zhenxi Guo. Part of structural computation was also performed on the Computing Platform of the Center for Life Science and High-performance Computing Platform of Peking University. We thank Yi Rao for sharing FLIPR instruments and Shangchen Han for technical supports. This work is supported by grants from the Ministry of Science and Technology of China (National Key R&D Program of China, 2016YFA0502004 to L.C.), National Natural Science Foundation of China (91957201, 31622021, 31870833 and 31821091 to L.C., 31900859 to J.-X. W.), Beijing Natural Science Foundation (5192009 to L.C.), Young Thousand Talents Program of China to L.C and the China Postdoctoral Science Foundation (2016M600856, 2017T100014, 2019M650324, and 2019T120014 to J.-X.W.). J.-X. W. is supported by the Boya Postdoctoral Fellowship of Peking University and the postdoctoral foundation of the Peking-Tsinghua Center for Life Sciences, Peking University (CLS).

## Author Contribution

Lei Chen initiated the project. Kangcheng Song, Miao Wei and Wenjun Guo purified proteins and prepared the cryo-EM samples. Kangcheng Song, Miao Wei, Wenjun Guo, Yunlu Kang and Jing-Xiang Wu collected the cryo-EM data. Kangcheng Song and Miao Wei processed the cryo-EM data, built and refined the atomic model with the help of Wenjun Guo and Lei Chen. Kangcheng Song performed the FLIPR experiments. All authors contributed to the manuscript preparation.

## Figure Legend

**Fig. S1.**
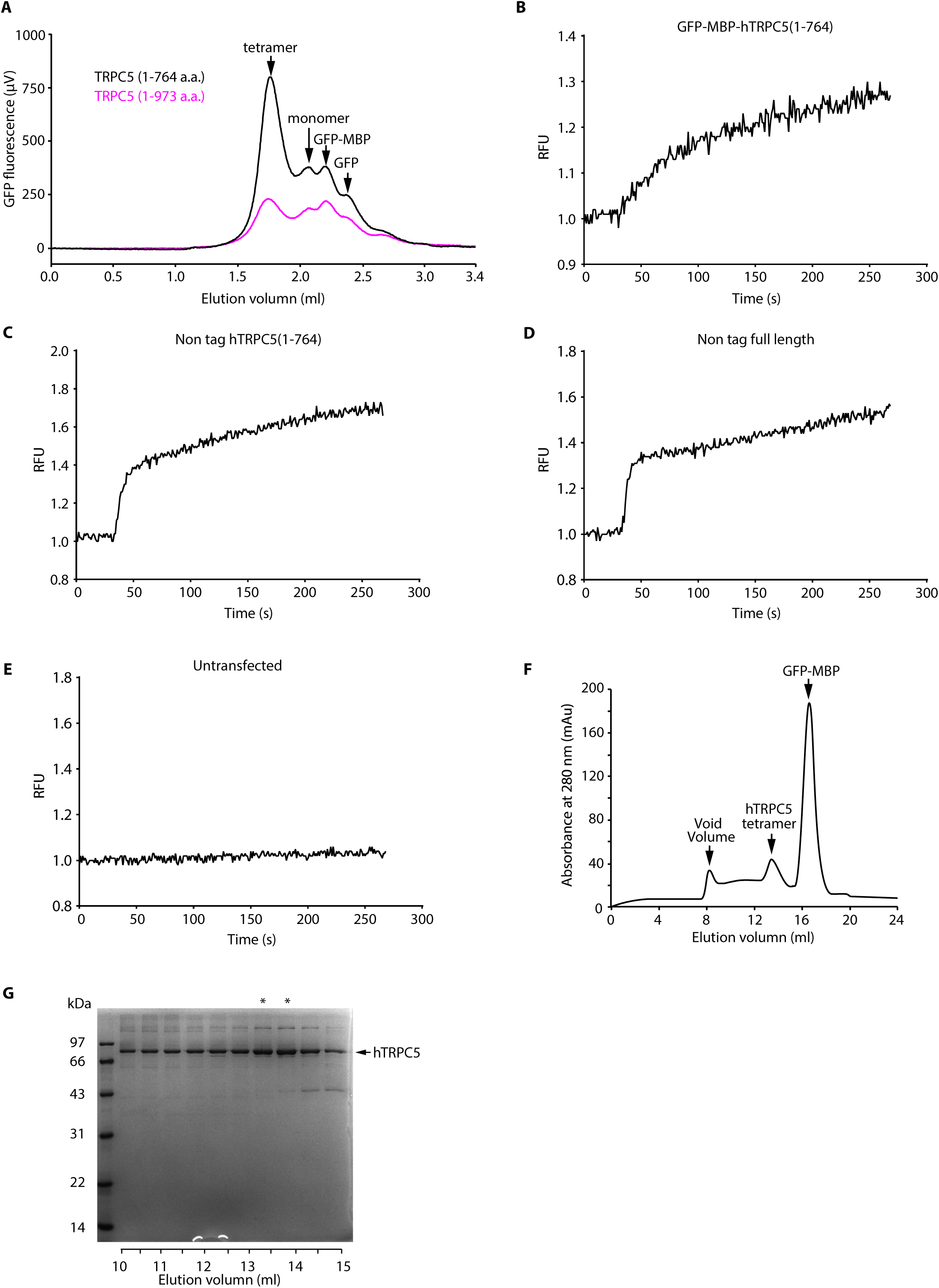
Functional characterization of full length hTRPC5 and truncated hTRPC5. (A) FSEC profile of full length hTRPC5 and hTRPC5_(1-764)_. (B-E) EA activation effect on PBM-NGFP-MBP-hTRPC5_(1-764)_, non-tag hTRPC5_(1-764)_, non-tag full length TRPC5 and untransfected cells detected by FLIPR. (F) Representative size-exclusion chromatography of hTRPC5 purified in GDN micelles. (G) SDS-PAGE gel of hTRPC5 eluted from size-exclusion chromatography. The position of TRPC5 is labeled and fractions that were pooled for cryo-EM analysis are denoted by asterisks.

**Fig. S2.**
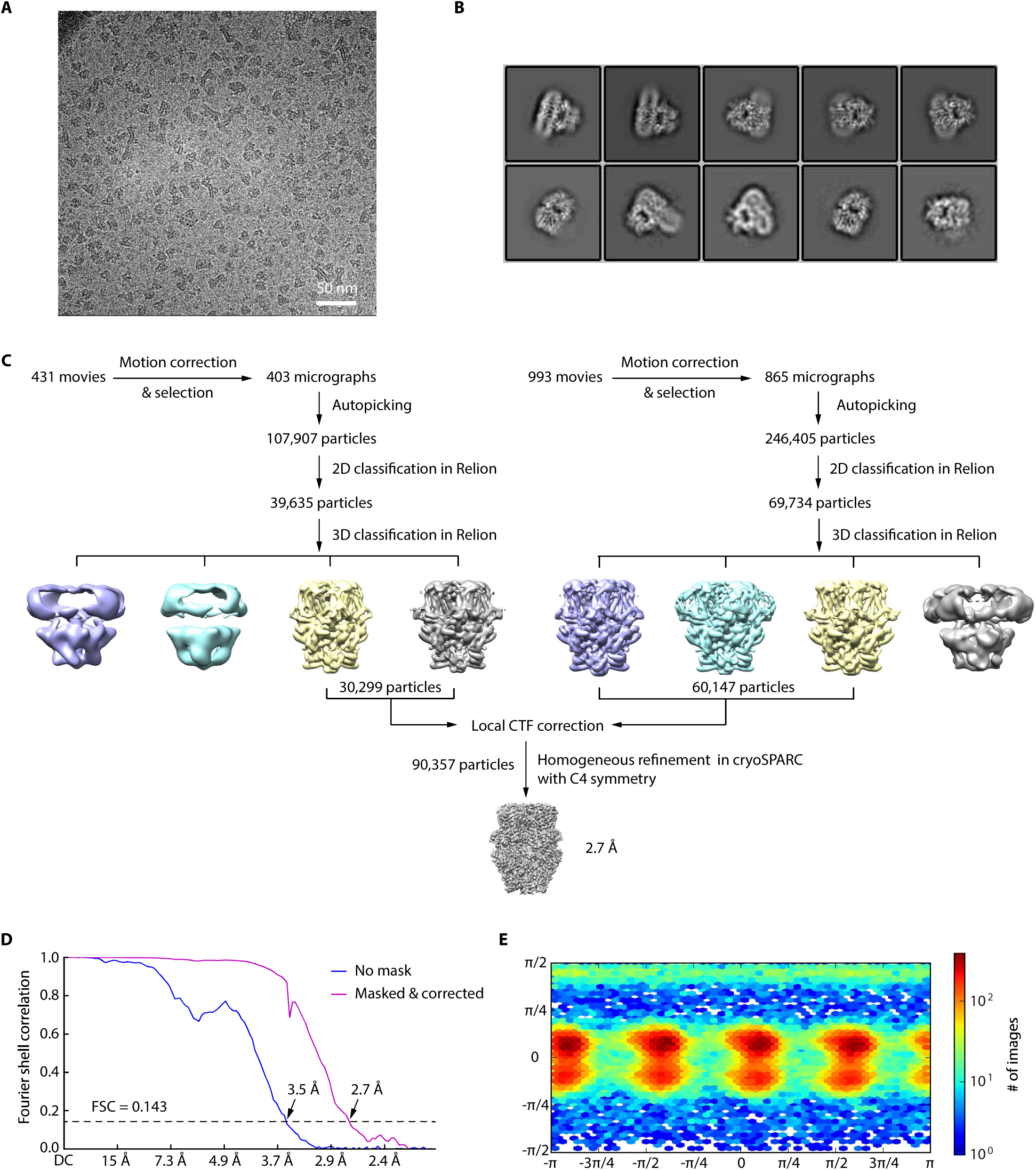
Cryo-EM image analysis of CMZ-bound hTRPC5. (A) Representative raw micrograph recorded on K2 Summit camera. (B) Representative 2D class averages of CMZ-bound hTRPC5. (C) Flowchart for Cryo-EM data processing of CMZ-bound hTRPC5. (D) Fourier shell correlation (FSC) curves of the two independently refined maps for unmasked (blue line, 3.5 Å) and corrected (purple line, 2.7 Å). Resolution estimation was based on the criterion of FSC 0.143 cut-off. (E) Angular distribution of CMZ-bound hTRPC5. This is a standard output from cryoSPARC.

**Fig. S3.**
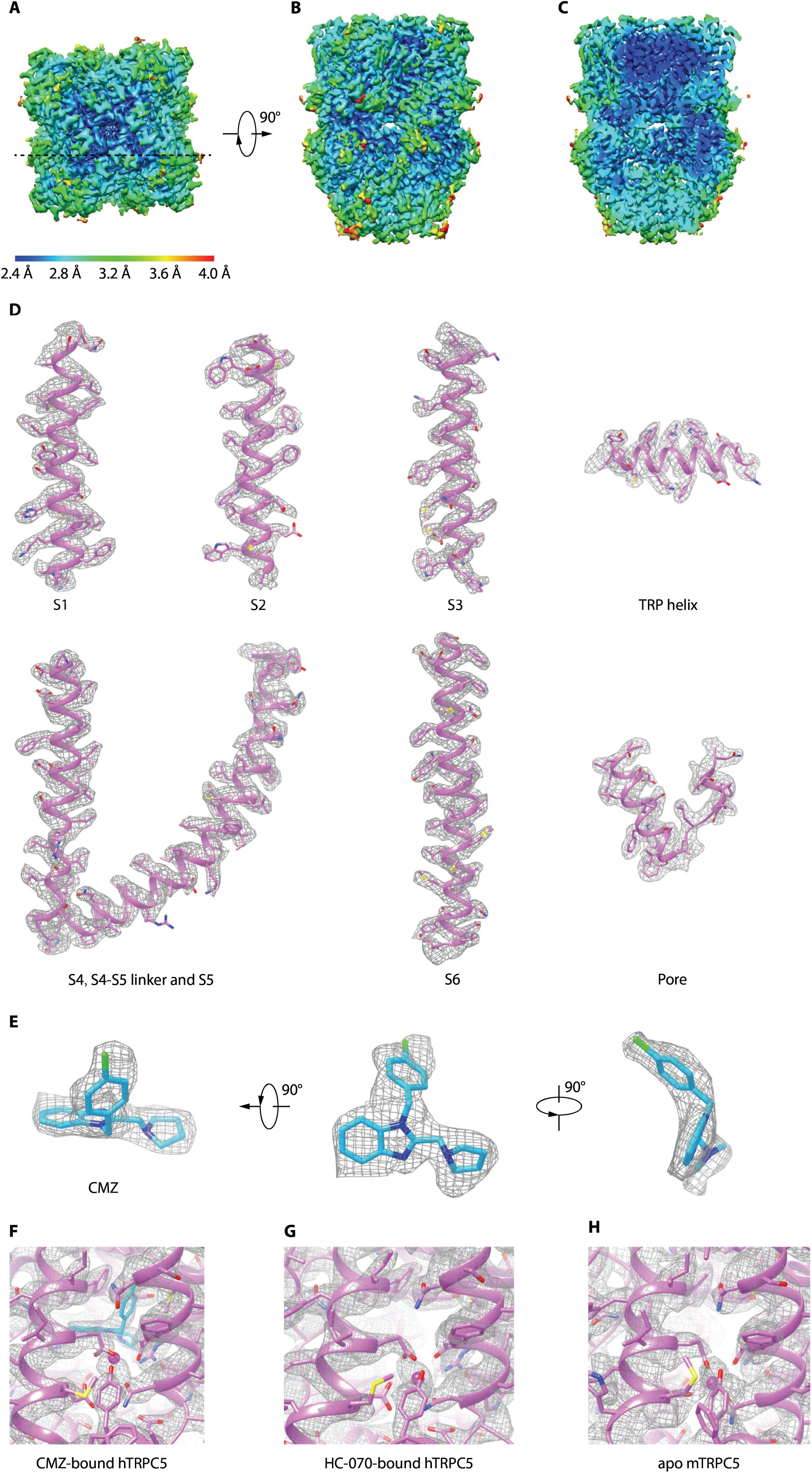
Cryo-EM map of CMZ-bound hTRPC5. (A-C) Cryo-EM map of CMZ-bound hTRPC5 colored by local resolution, shown in top view (A), side view (B), and cross-section (C). The position of cross-section is shown as dashed line in (A). (D) Cryo-EM density map (contoured at 3.6σ, gray mesh) with atomic models superimposed. (E) Cryo-EM density of CMZ shown in different views (contoured at 3.5σ). (F) Cryo-EM densities of CMZ-binding pocket in CMZ-bound TRPC5 map (contoured at 5σ). (G) Cryo-EM densities in HC-070-bound TRPC5 and apo mTRPC5 maps corresponding to the CMZ-binding pocket contoured at the same σ as in (F).

**Fig. S4.**
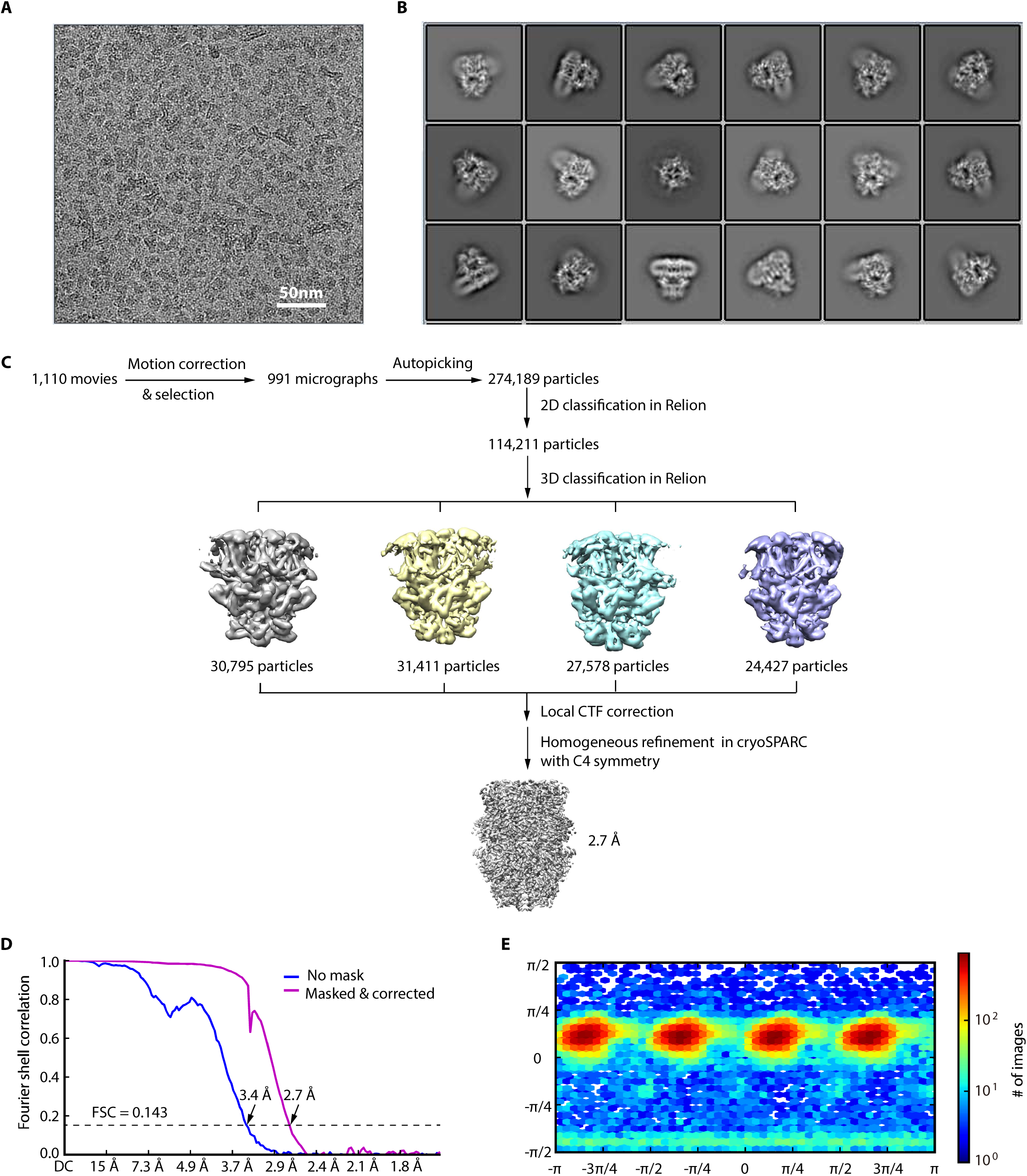
Cryo-EM image analysis of HC-070-bound hTRPC5. (A) Representative raw micrograph recorded on K2 Summit camera. (B) Representative 2D class averages of HC-070-bound hTRPC5. (C) Flowchart for cryo-EM data processing of HC-070-bound hTRPC5. (D) Fourier shell correlation (FSC) curves of the two independently refined maps for unmasked (blue line, 3.4 Å) and corrected (purple line, 2.7 Å). Resolution estimation was based on the criterion of FSC 0.143 cut-off. (E) Angular distribution of HC-070-bound hTRPC5. This is a standard output from cryoSPARC.

**Fig. S5.**
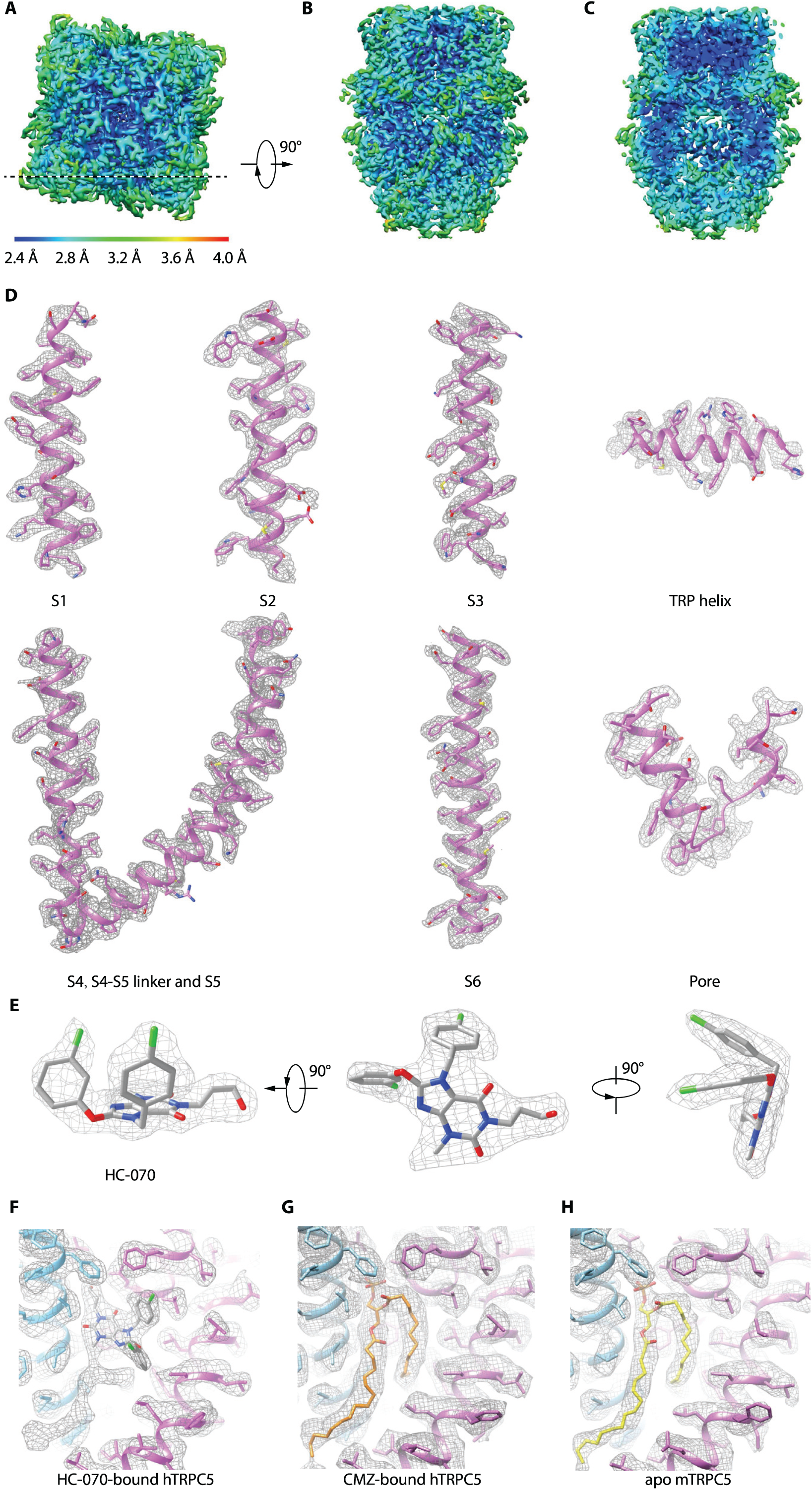
Cryo-EM map of HC-070-bound hTRPC5. (A-C) Cryo-EM map of HC-070-bound hTRPC5 colored by local resolution, shown in top view (A), side view (B) and cross-section (C). The position of cross-section is shown as dashed line in (A). (D) Cryo-EM density map (contoured at 3σ, gray mesh) with atomic model superimposed. (E) Cryo-EM density map of HC-070 shown in different views (contoured at 3σ). (F-H) Cryo-EM densities of HC-070-binding pocket in (F) HC-070-bound hTRPC5 map (contoured at 3.6σ), (G) CMZ-bound hTRPC5 map (contoured at 2σ), and (H) apo mTRPC5 map (contoured at 3.1σ).

**Fig. S6.**
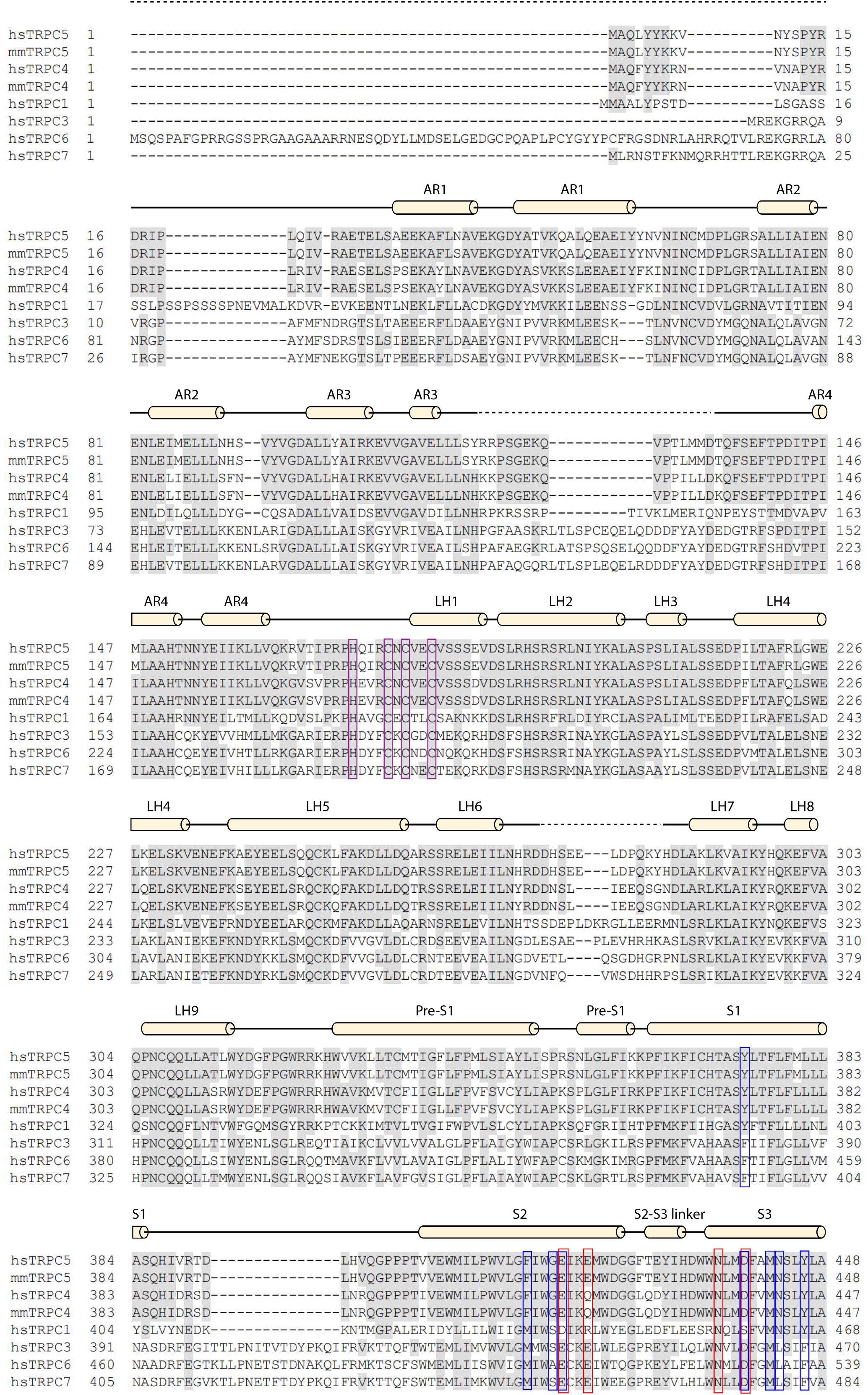
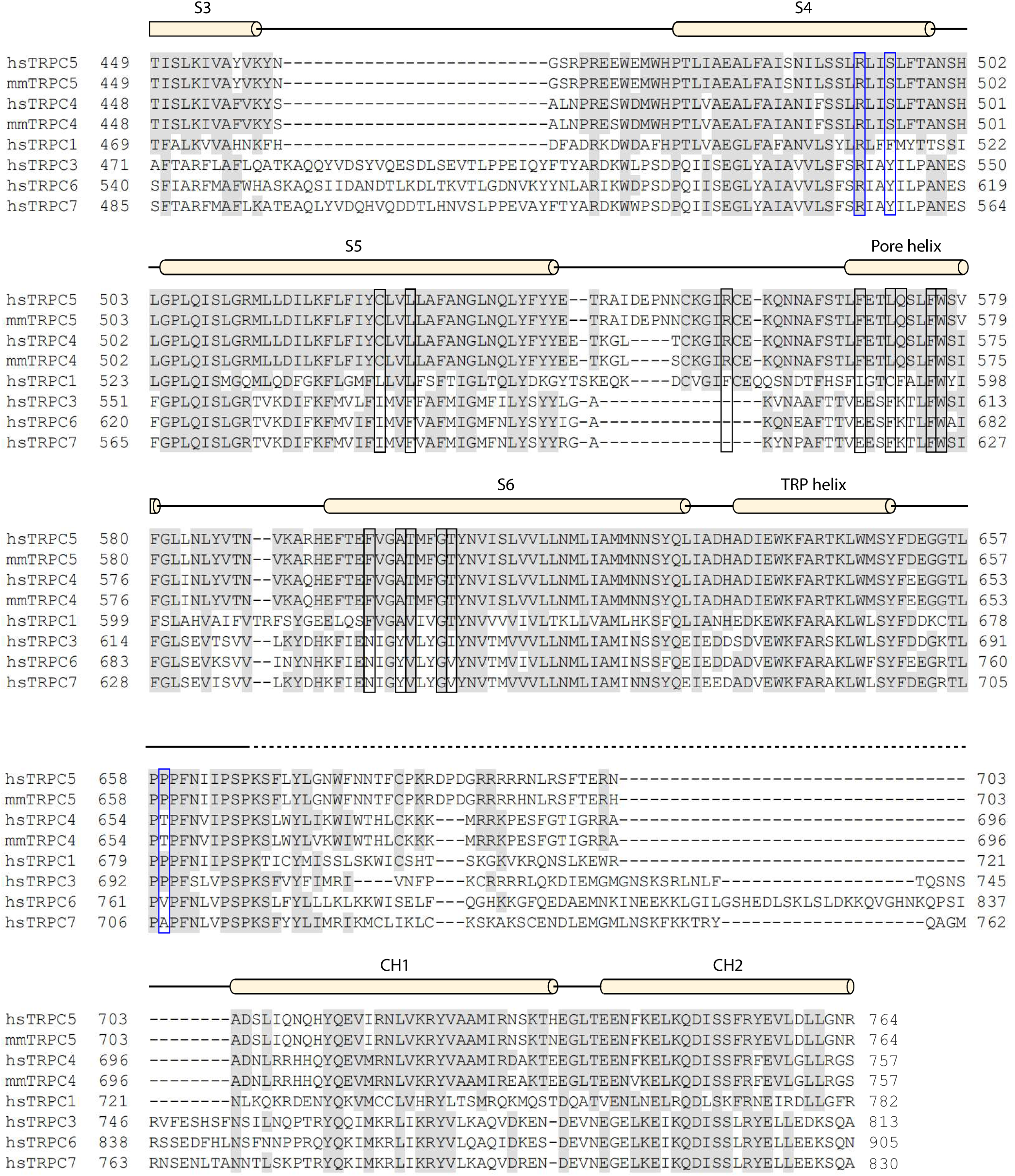
Sequence alignment of the TRPC channels. Conserved residues are colored in gray. Secondary structures are indicated as cylinders (α helices) and lines (loops). Unmodeled residues are indicated as dashed line. Residues of CMZ binding site, HC-070 binding site, VSLD’s cation binding site and zinc binding site in CTD are indicated with blue boxes, black boxes, red boxes and purple boxes respectively.

**Fig. S7.**
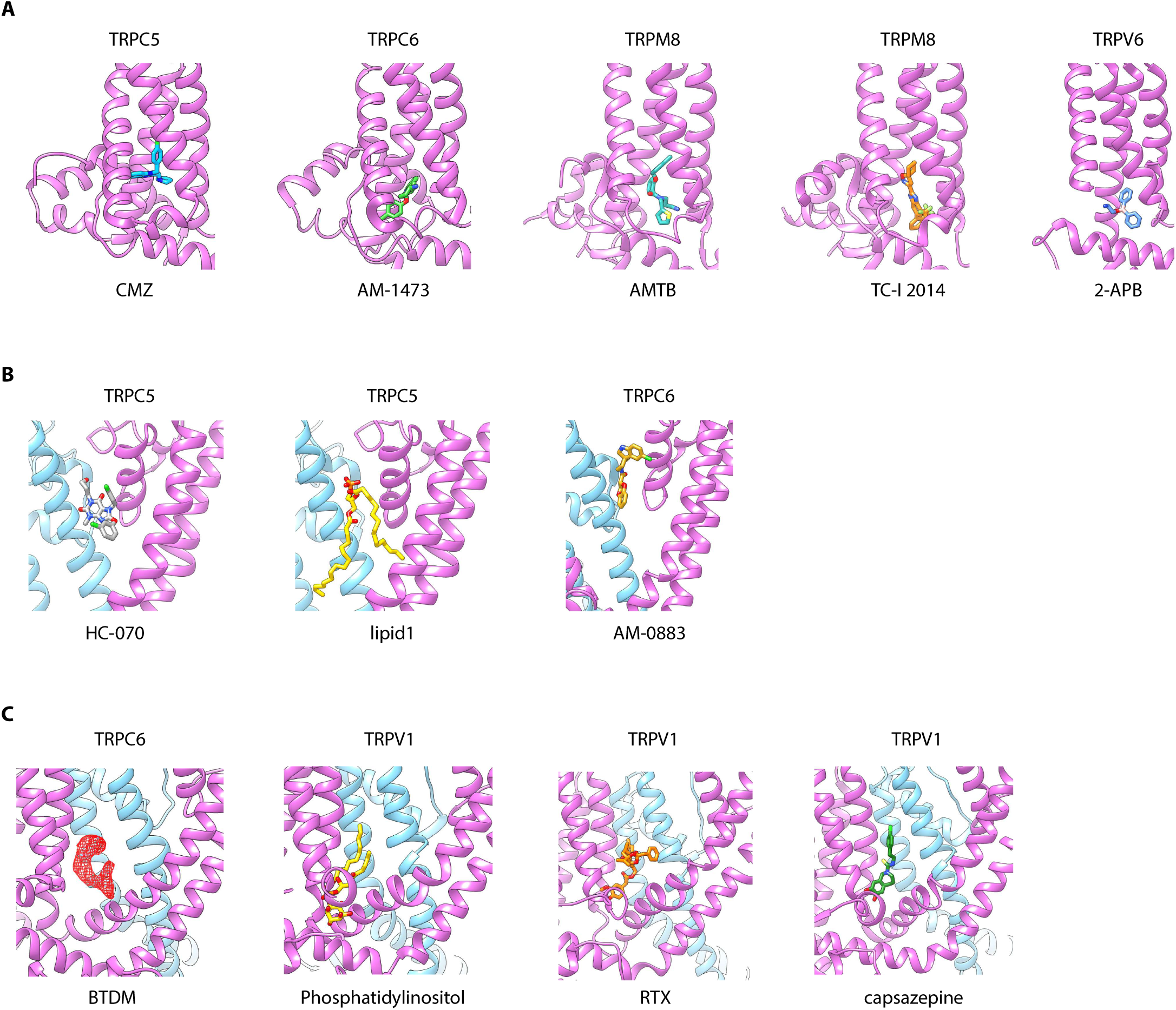
Ligand binding sites in TRP family. (A) Different ligand binding in IBP-A among TRPC5, TRPC6, TRPM8 and TRPV6. One subunit of different TRPCs are shown in cartoon and colored in purple. Different ligands are shown as sticks and in various color. (B) Different ligand binding in IBP-B among TRPCs. One subunit and the adjacent subunit are colored in purple and light blue, respectively. (C) Different ligand binding in IBP-C among TRPC6 and TRPV1. The density of BTDM is shown in mesh and other modeled ligands are shown as sticks.

**Fig. S8.**
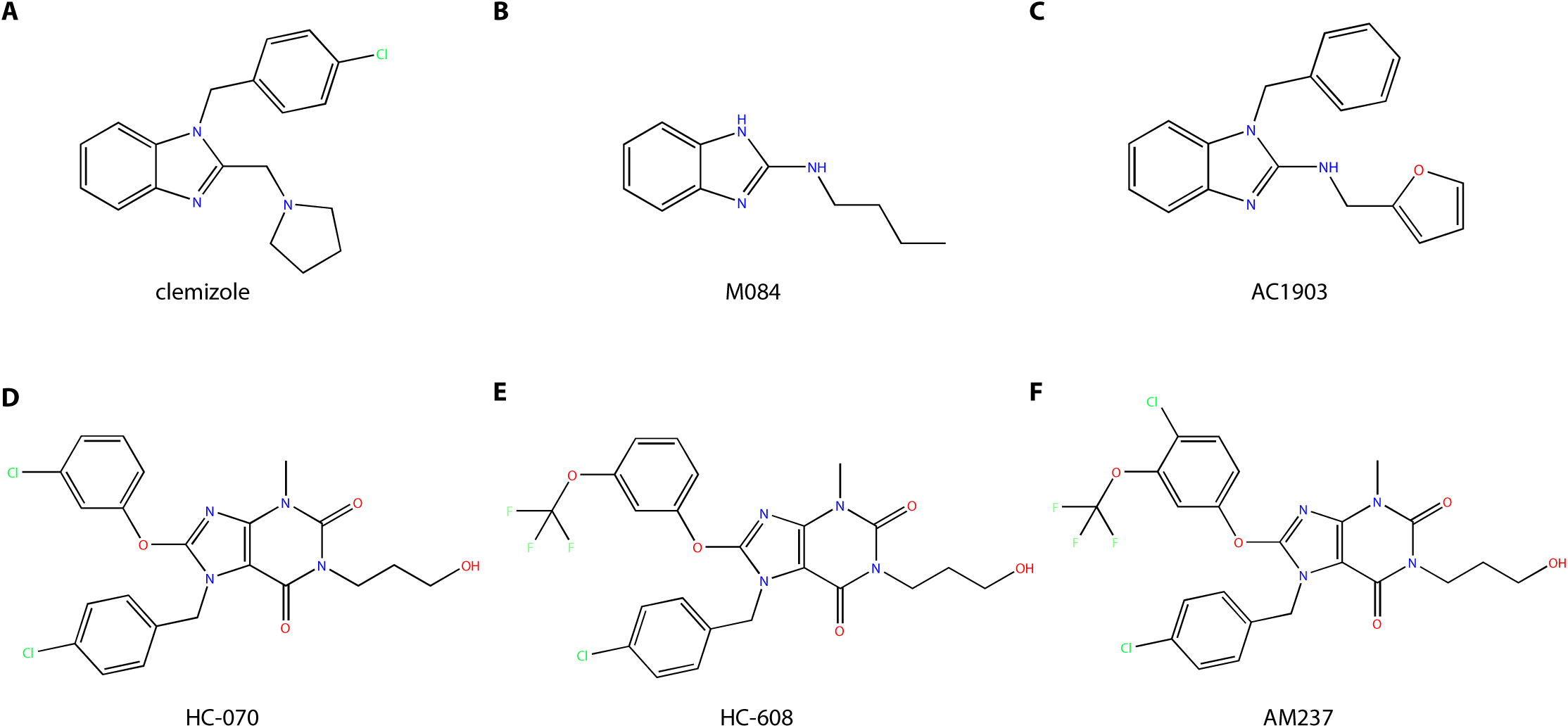
Chemical structure comparison between different TRPC5 ligands. (A-C) Chemical structure comparison between benzimidazole derivatives inhibitors: CMZ (A), M084 (B) and AC1903 (C). (D-F) Chemical structure comparison between xanthines derivatives ligands: inhibitors HC-070 (D), and HC-608 (E) and activator AM237 (F).

**Supplementary Table. 1.** Cryo-EM data collection, refinement and validation statistics

| PDB ID<br>EMDB ID | CMZ-bound<br>hTRPC5<br>XXXX<br>EMD-XXXX | HC-070-bound<br>hTRPC5<br>XXXX<br>EMD-XXXX |
| --- | --- | --- |
| **Data collection and processing** |  |  |
| Magnification | 130,000 × | 165,000 × |
| Voltage (kV) | 300 | 300 |
| Electron exposure (e-/Å<sup>2</sup>) | 50 | 50 |
| Defocus range (μm) | -1.5 to -2.0 | -1.5 to -2.0 |
| Pixel size (Å) | 1.045 | 0.821 |
| Symmetry imposed | *C4* | *C4* |
| Initial particle images (no.) | 354,312 | 274,189 |
| Final particle images (no.) | 90,446 | 114,211 |
| Map resolution (Å) | 2.74 | 2.74 |
| FSC threshold | 0.143 | 0.143 |
| Map resolution range (Å) | 250-2.7 | 250-2.7 |
| **Refinement** |  |  |
| Initial model used (PDB code) | 5Z96 | 5Z96 |
| Model resolution (Å) | 250-2.7 | 250-2.7 |
| FSC threshold | 0.143 | 0.143 |
| Model resolution range (Å) | 250-2.7 | 250-2.7 |
| Map sharpening *B* factor (Å<sup>2</sup>) | -119.0 | -118.1 |
| Model composition |  |  |
| Non-hydrogen atoms | 23,200 | 23,256 |
| Protein residues | 2,756 | 2,776 |
| Ligands | 28 | 24 |
| *B* factors (Å<sup>2</sup>) |  |  |
| Protein | 117.47 | 130.08 |
| Ligand | 104.17 | 40.16 |
| R.m.s. deviations |  |  |
| Bond lengths (Å) | 0.005 | 0.013 |
| Bond angles (°) | 1.035 | 1.402 |
| Validation |  |  |
| MolProbity score | 2.02 | 2.94 |
| Clashscore | 8.38 | 29.62 |
| Poor rotamers (%) | 3.58 | 7.42 |
| Ramachandran plot |  |  |
| Favored (%) | 97.21 | 95.34 |
| Allowed (%) | 2.79 | 4.66 |
| Disallowed (%) | 0.00 | 0.00 |

